# Age Associated Shifts Toward Anaerobic Taxa and Amino Acid Degradation Potential in the Canine Oral Microbiome

**DOI:** 10.64898/2026.10.30.763000

**Authors:** Zhiyue Jiang, Madeline Shono, Aisling Archdeacon, Giorgia Del Vecchio, Matteo Pellegrini

**Affiliations:** UCLA Department of Molecular, Cell and Developmental Biology

**Keywords:** canine oral microbiome, aging, shotgun metagenomics, anaerobic bacteria, microbial metabolism, periodontal disease

## Abstract

**Introduction:** Periodontal disease is common in dogs and is associated with changes in oral microbial community composition. How the canine oral microbiome restructures across age at both ecological and metabolic levels remains incompletely characterized.

**Methods:** We analyzed shotgun metagenomic buccal-swab profiles from 1,125 dogs using MetaPhlAn. Functional profiles from HUMAnN were analyzed at pathway and functional-group levels. We annotated species-level taxa with oxygen preference and evaluated oxygen-tolerant and anaerobic taxa abundances across age. We also characterized community structure, metabolic potential, Shannon diversity, and age and weight prediction using nested cross-validated Elastic Net models.

**Results:** Age was associated with substantial restructuring in the oral microbiome. Mean oxygen-tolerant taxa abundance decreased from 67.5% in puppies to 29.1% in mature dogs, whereas anaerobic abundance increased from 14.3% to 53.8%. The oxygen-tolerant: anaerobe log-ratio declined with age (Spearman ρ = −0.616). Metabolic profiles also varied with age, with higher amino-acid-degradation potential and lower glycolysis and central-carbohydrate potential, and aerobic-respiration-I abundance in older dogs. Shannon diversity increased with age (ρ = 0.537, p < 0.001). Elastic Net models trained on microbiome species abundance predicted age (Pearson r = 0.609; R^2^ = 0.371) and weight (r = 0.498; R^2^ = 0.248) with moderate accuracy.

**Discussion:** Canine age was associated with coordinated changes in oral-community oxygen preference and metabolic pathways. These findings extend prior taxon-specific observations by examining a broader ecological shift and metabolic potential in the canine oral microbiome. This provides a reference for future longitudinal or clinical studies of oral maturation and periodontal disease.

## 1. Introduction

Dogs are a valuable model system for biological studies because of their phenotypic diversity. Despite belonging to a single species, dogs show substantial variation in morphology and body size, making them useful for investigating host factors associated with microbial communities. Dogs also naturally develop age-associated conditions, including periodontal disease. Their oral microbiome contains taxa related to those implicated in human periodontal disease, although the two host species do not have identical oral microbial ecologies.

Periodontal disease (PD) is among the most prevalent oral diseases in dogs. PD affects up to 80% of adult canines above 3 years old (Enlund et al., 2020). It may go unrecognized during early development as comprehensive periodontal assessment generally requires examination under anesthesia, which can complicate timely diagnosis and treatment (Niemiec, 2008; Wallis and Holcombe, 2020). The disease exists as a spectrum from gingivitis to periodontitis. Gingivitis is a reversible inflammatory response of the gums, whilst periodontitis causes irreversible destruction of connective tissue and tooth loss.

Rather than being caused by a single pathogen, disease progression is accompanied by large-scale restructuring of the oral microbiome, including shifts in oxygen utilization, metabolic activity, and community composition (Niemiec et al., 2022). Dogs with periodontal disease often have greater relative abundance of anaerobe-associated taxa, including *Porphyromonas, Treponema*, and *Tannerella*, whereas several aerobe-associated taxa are more common in clinically healthy oral communities (Davis et al., 2013; Santibanez et al., 2021; Niemiec et al., 2022). In humans, Porphyromonas, Treponema, and Tannerella are also associated with periodontal disease. Related groups have also been reported in canine disease, although species-specific roles can differ (Di Bello et al., 2014; Davis et al., 2013; Santibáñez et al., 2021). These observations motivate community-level study of age-associated microbiome changes.

Age is a major risk factor for canine periodontal disease, with prevalence increasing substantially throughout the lifespan (O’Neill et al., 2021). Body size is also associated with oral health, with smaller dogs generally exhibiting higher rates of periodontal disease (O’Neill et al., 2021). However, cross-sectional microbiome studies without periodontal examinations cannot determine whether a taxon changes because of normal maturation, accumulated plaque, undiagnosed disease, breed-related factors, or other elements. Age-associated microbial changes are thus potential indicators of oral-health risk, but cannot replace periodontal examinations.

Previous work has reported age-associated changes in canine oral microbial diversity and in taxa such as Conchiformibius steedae and Porphyromonas (Templeton et al., 2023; Kislik et al., 2024). Here, we examined age- and weight-associated variation at sample, order, and genus levels and tested whether microbial composition could predict host age and weight. We used HUMAnN to test whether age-associated taxonomic restructuring was accompanied by shifts in metabolic pathways. We hypothesized that taxa associated with oxygen-tolerant oral communities would be more abundant in younger dogs, whereas taxa previously reported in canine periodontal-disease studies would be more abundant in older dogs. Because clinical periodontal status was unavailable, we interpret these patterns as age-associated differences in microbiome composition rather than evidence of disease in individual dogs.

## 2. Methods

### 1. DNA extraction and whole-genome library preparation

Canine buccal swabs were sequenced via whole-genome shotgun sequencing. Canine DNA was collected using an animal-specific saliva collection kit (PERFORMAgene PG-100, DNA Genotek). Following sample collection, tubes were incubated overnight at 50°C. Genomic DNA (gDNA) was extracted by adding 0.65× volumes of nuclease-free water and 0.65× volumes of SPRI magnetic beads (Beckman Coulter). After thorough mixing, samples were incubated at room temperature for 5 minutes and then placed on a magnetic rack for an additional 5 minutes to separate the beads. The beads were washed twice with 80% ethanol, air-dried for 5 minutes, and the purified gDNA was eluted in EB buffer (Qiagen). DNA concentration was determined using the Qubit dsDNA BR Assay Kit (Thermo Fisher Scientific). Whole-genome sequencing libraries were prepared from 150 ng of gDNA using the xGen DNA Library Preparation Kit (Integrated DNA Technologies). Briefly, genomic DNA underwent enzymatic fragmentation followed by end repair and A-tailing. Full-length IDT Y-shaped UDI+UMI adapters were then ligated to the DNA fragments. Following ligation, eight samples were pooled and purified using a 0.8× SPRI bead cleanup. DNA was eluted in 20 µL of EB buffer, and 10 µL was used for PCR amplification (1 cycle). Final libraries were quality-controlled using the Qubit dsDNA BR Assay Kit (Thermo Fisher Scientific) and the Agilent TapeStation D1000 assay. Libraries were sequenced on an Illumina NovaSeq X Plus platform using 2 × 100 bp paired-end reads.

### 2. Taxonomic profiling

Forward reads (R1) were profiled with MetaPhlAn version 4.1.1 using the mpa_vJan25_CHOCOPhlAnSGB_202503 database bundle. MetaPhlAn estimates relative abundance using clade-specific marker genes and Bowtie2 alignments (Blanco-Míguez et al., 2023). Relative abundances were restricted to the bacterial kingdom and normalized such that total bacterial abundance summed to 1 within each sample. For abundance-based multivariate analyses, abundances were transformed as log_10_(*abundance* + 1 × 10^−5^) to stabilize variance and reduce skewness.

### 3. Quality control and analytical cohort

Taxonomic profiles were available for 1,125 dogs. PCA followed by UMAP and DBSCAN (as described below) identified an external cluster of 80 dogs with a mean Enterobacterales relative abundance of 58.35%. These 80 dogs were excluded from downstream analyses, leaving 1,045 dogs. Analyses involving weight excluded dogs younger than 1 year to reduce confounding by developmental growth.

### 4. Order-level community composition

Order-level abundance profiles were clustered using SciPy (scipy.cluster.hierarchy.linkage) using correlation distance with average linkage to identify groups of taxa and samples with similar abundance patterns. Samples were annotated by host age.

### 5. UMAP and pseudotime

UMAP and pseudotime analyses were performed in Python using scikit-learn, umap-learn, and SciPy (Pedregosa et al., 2011; McInnes et al., 2018; Virtanen et al., 2020). PCA was performed on normalized and log-transformed order-level abundance profiles, and UMAP was generated from the retained principal components. DBSCAN implemented in scikit-learn was used to identify the external Enterobacterales-enriched cluster. Pseudotime was calculated from a nearest-neighbor graph constructed in PCA space, using the youngest dog (DW31240411301134; approximately one week old) as the root.

A separate UMAP was constructed with umap-learn to visualize relationships among bacterial orders. Five orders with no abundance variation across dogs were excluded, leaving 46 orders. We used correlation distance, 12 nearest neighbors, a minimum distance of 0.25, and random state 42. Each point represents a bacterial order; point color indicates its Spearman correlation with host age, and point size represents its mean relative abundance.

### 6. Nonlinear genus trends

We evaluated associations between host age and the abundance of every genus using SciPy 1.17.1 and adjusted p-values by the Benjamin-Hochberg procedure with statsmodels 0.14.6. Nonlinear trends were visualized with LOWESS using statsmodels.nonparametric.smoothers_lowess.lowess (smoothing fraction = 0.65).

### 7. Alpha diversity analysis

Shannon diversity indices (H) were calculated at the species level to quantify alpha diversity across samples. The index was implemented directly with NumPy 2.4.4 using natural logarithms. Diversity trends were evaluated continuously against age (years) and compared across age bins (0–2, >2–7, and >7 years). Similarly, diversity trends were evaluated amongst weight groups (<20, 20–<60, and ≥60 lbs). The formula for Shannon diversity indices is: 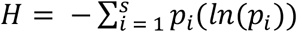 where p is the relative abundance of the ith species within a dog, s is the total number of species, and i is the ith species. Corresponding Spearman correlation coefficients were calculated with scipy.stats.spearmanr.

### 8. Species oxygen preference annotation

Bacterial species were annotated using evidence from primary literature and BacDive (Schober et al., 2025). Phenotypes were labeled as obligate aerobe, aerobe, facultative anaerobe, microaerophile, variable oxygen-tolerant, anaerobe, or unknown. For downstream analysis, the first five categories were grouped as oxygen-tolerant and compared with anaerobes. For each dog, we calculated 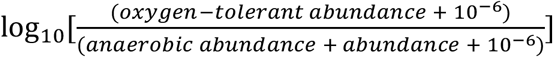 and assessed its association with age using Spearman correlation. The numerator and denominator are summed relative abundances of species assigned to the respective groups. Positive log-ratios indicate greater oxygen-tolerant abundance, whereas negative values indicate greater anaerobic abundance.

### 9. Functional pathway profiling (HUMAnN 3)

Paired-end reads were cleaned with KneadData version 0.10.0, which used the ROS_Cfam_1.0 dog reference genome with default parameters. Remaining plus orphaned reads were concatenated for each sample and functionally profiled with HUMAnN version 3.9 (Beghini et al., 2021). Functional profiling utilized three databases: MetaPhlAn species marker gene database mpa_vJun23_CHOCOPhlAnSGB_202403, UniRef90 for proteins, and ChocoPhlAn for nucleotides. Sample outputs were joined together, and community pathway abundance tables were normalized to copies per million (CPM), excluding UNMAPPED and UNINTEGRATED pathways. Samples with at least 2 million reads post-cleaning were retained (n = 623).

A pathway was considered detected when its HUMAnN abundance was greater than zero CPM. Pathways detected in at least 10% of dogs were retained for downstream analyses (n = 301). For each dog, centered log-ratio (CLR) values were calculated as ln(*CPM* + 1)for each retained pathway minus that dog’s mean ln(*CPM* + 1)across all retained pathways. CLR-transformed pathway profiles were analyzed by PCA using scikit-learn. Spearman correlations tested associations between age, the number of detected pathways, PC1, and PC2. Complete PC1 and PC2 loadings are provided in Supplementary Table S1.

Pathways were assigned to functional groups using the MetaCyc_pathway_map dataset from the R package file2meco. This reference organizes MetaCyc pathways into a hierarchical classification containing two levels of functional superclasses and individual pathways. Pathways were matched to this reference to group related metabolic functions into broader categories. Functional-group scores summarized standardized log-transformed pathway abundances and represented functional potential. Age associations were evaluated using Spearman correlation with Benjamini–Hochberg correction.

### 10. Multivariate age models

Elastic Net regression (scikit-learn, sklearn.linear_model.ElasticNet) was used to evaluate whether oral microbiome composition predicted host age or body weight. The age-model cohort was obtained by excluding the 369 dogs younger than one year and 57 that lacked weight information. Age models included 676 dogs at least 1 year old; weight models included 619 of these dogs with recorded weight. StandardScaler and ElasticNet were combined in a scikit-learn Pipeline so that scaling parameters were estimated from each training fold. Hyperparameters were selected with GridSearchCV and shuffled KFold splits. Separate models were fit for age and body weight using nested cross-validation. In each of 10 shuffled outer folds (random state = 42), hyperparameters were selected within the training data using 5-fold shuffled inner cross-validation and negative mean squared error. Alpha was searched over [1 × 10^−5^, 1 × 10^−4^, 1 × 10^−3^, 1 × 10^−2^, 0.1, 1, 10, 100], and the L1/L2 mixing parameter was searched over [0.2, 0.4, 0.6, 0.8].

The model selected within each outer-fold training set was used to generate a prediction for its held-out samples. Performance was evaluated from pooled held-out predictions using Pearson correlation and its associated p-value, R^2^, mean squared error, root mean squared error, mean absolute error, and median absolute error.

## 3. Results

### 1. Unsupervised clustering reveals restructuring of the microbiome with age

We analyzed whole-genome shotgun sequencing profiles generated from buccal swabs collected from 1,125 dogs. MetaPhlAn profiles were aggregated to bacterial orders and clustered with correlation distance and average linkage (see Methods). The resulting clustermap showed structured variation across samples, with bacterial orders grouping by similar abundance patterns (Figure 1).

**Figure 1.**
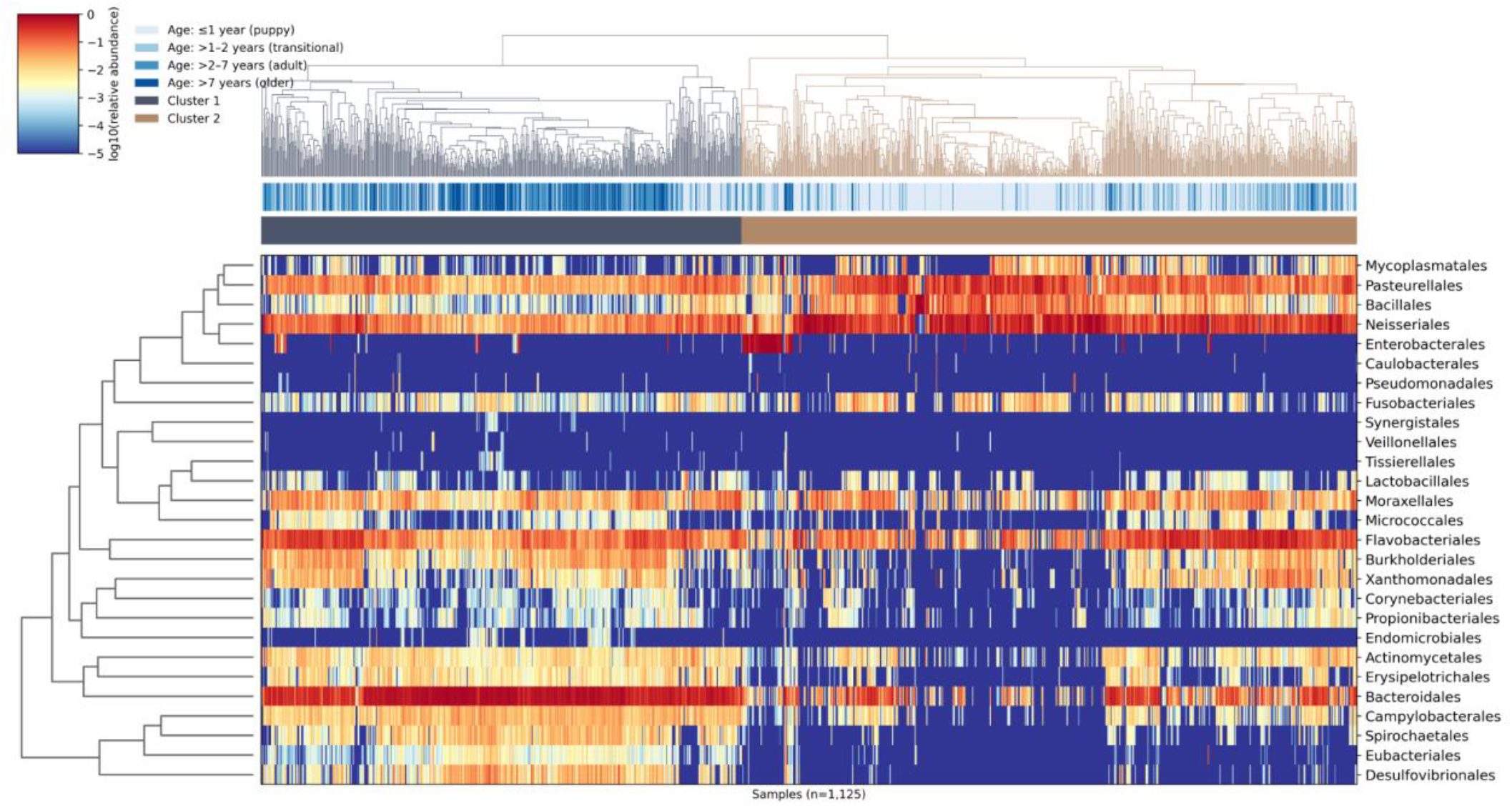
Unsupervised order-level clustermap of canine oral microbiome composition. Columns represent individual dogs, and rows represent bacterial orders. Upper annotation strip indicates dog age group. Dendrogram branch colors and lower annotation strip indicate the two sample clusters.

Although age was not used to generate the clustering, sample age annotations showed age dependence across the heatmap. Hierarchical clustering separated the samples into two major community-composition clusters (Figure 1). One cluster contained 493 dogs and was dominated by Bacteroidales, with a mean relative abundance of 52.4%. The second contained 632 dogs and had greater Neisseriales and Pasteurellales abundance. Mean Bacteroidales abundance was 9.0% in this cluster, while Neisseriales and Pasteurellales averaged 33.4% and 16.1%. Age was not used to define the clusters, but the Bacteroidales-rich cluster contained generally older dogs, with a median age of 4.0 years compared with 0.83 years in the Neisseriales/Pasteurellales-rich cluster.

### 2. Sample-level UMAP shows a continuous age-dependent microbiome gradient

To further characterize microbiome community structure, we projected the 1,125-dog dataset onto two dimensions using UMAP (Figure 2). Age varied across the ordination, consistent with differences in community composition across dog ages. Coloring the same ordination by the oxygen-tolerant: anaerobe log-ratio (see Methods: Species oxygen preference annotation) showed that this physiological balance varied along the community gradient (Figure 2B). A distinct 80-dog cluster, highlighted in magenta, had a mean Enterobacterales relative abundance of 58.35%, compared with 0.010% among the remaining dogs. This cluster was excluded from downstream analyses.

**Figure 2.**
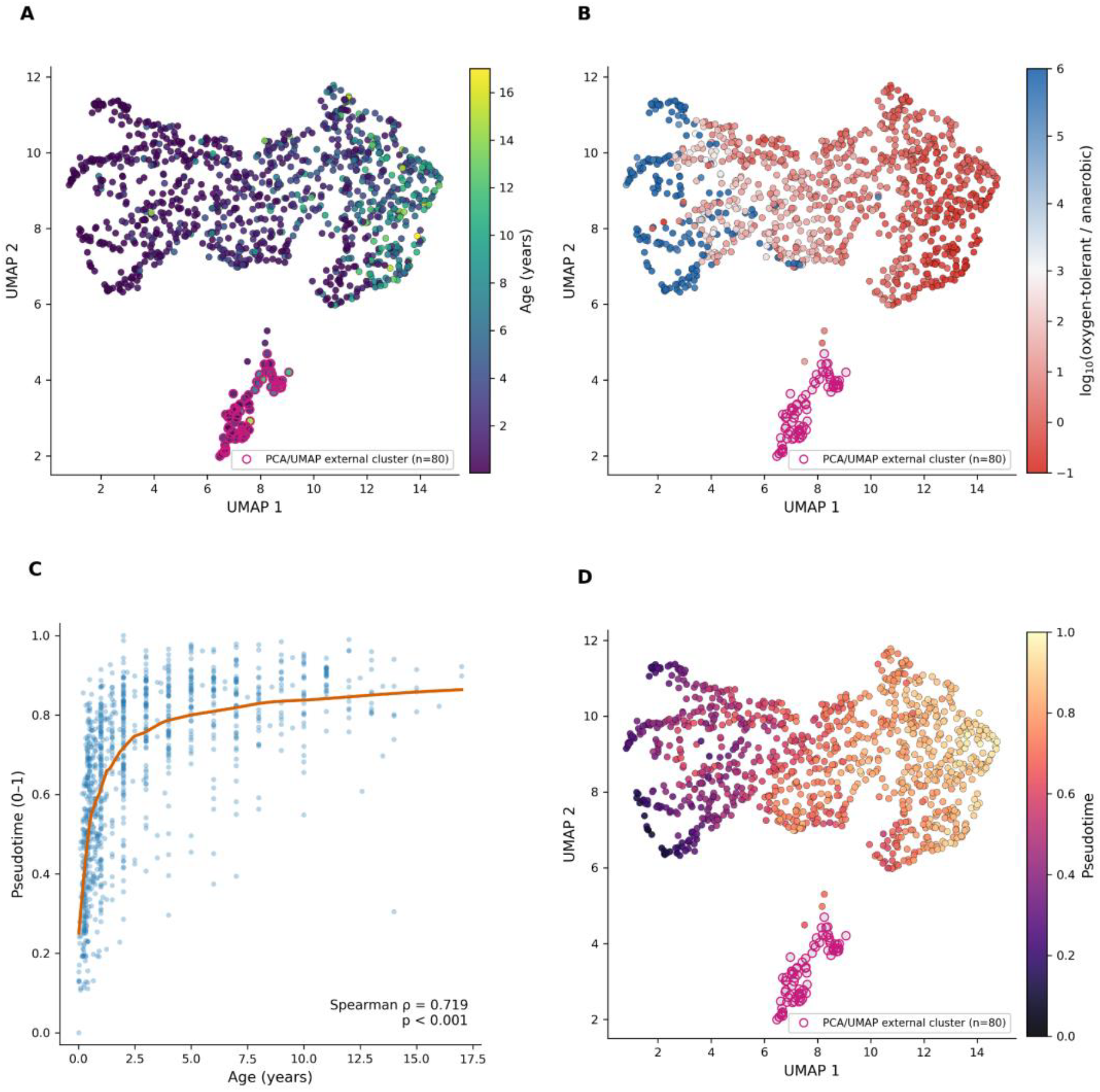
Continuous age-associated trajectory of the canine oral microbiome. (A) UMAP projection of dog microbiome samples colored by host age. (B) The same UMAP colored by the log10 ratio of oxygen-tolerant to anaerobic relative abundance, using a scale from −1 to 6. (C) Relationship between host age and UMAP-derived pseudotime. (D) Dog UMAP colored by pseudotime. The 80 dogs in the external Enterobacterales-enriched cluster are highlighted with identical magenta point outlines in panels A, B, and D.

Pseudotime was calculated using a nearest-neighbor graph constructed in PCA space (Figure 2C). The 80 samples in a disconnected component were not assigned pseudotime because no graph path connected them to the root dog (see Methods). Pseudotime was strongly associated with age (Spearman ρ = 0.719, p < 0.001) but the LOWESS trend was nonlinear. The largest change occurred during the first year of life and the trajectory was more gradual at older ages. Mapping pseudotime onto the UMAP visualized the same cross-sectional community gradient (Figure 2D).

### 3. Bacterial-level analyses identify taxa associated with age

A UMAP of bacterial order was used to visualize how taxa with different age associations were distributed in the taxonomic feature space (Figure 3A). Bacteroidales included anaerobic species such as *Porphyromonas gulae, Porphyromonas cangingivalis*, and *Tannerella forsythia*, which increased with age. Neisseriales included oxygen-tolerant species such as *Conchiformibius steedae* and *Neisseria animaloris*, while Pasteurellales included the microaerophile *Canicola haemoglobinophilus* and facultative anaerobe *Frederiksenia canicola*. This pattern aligned with the broader oxygen-preference analysis.

**Figure 3.**
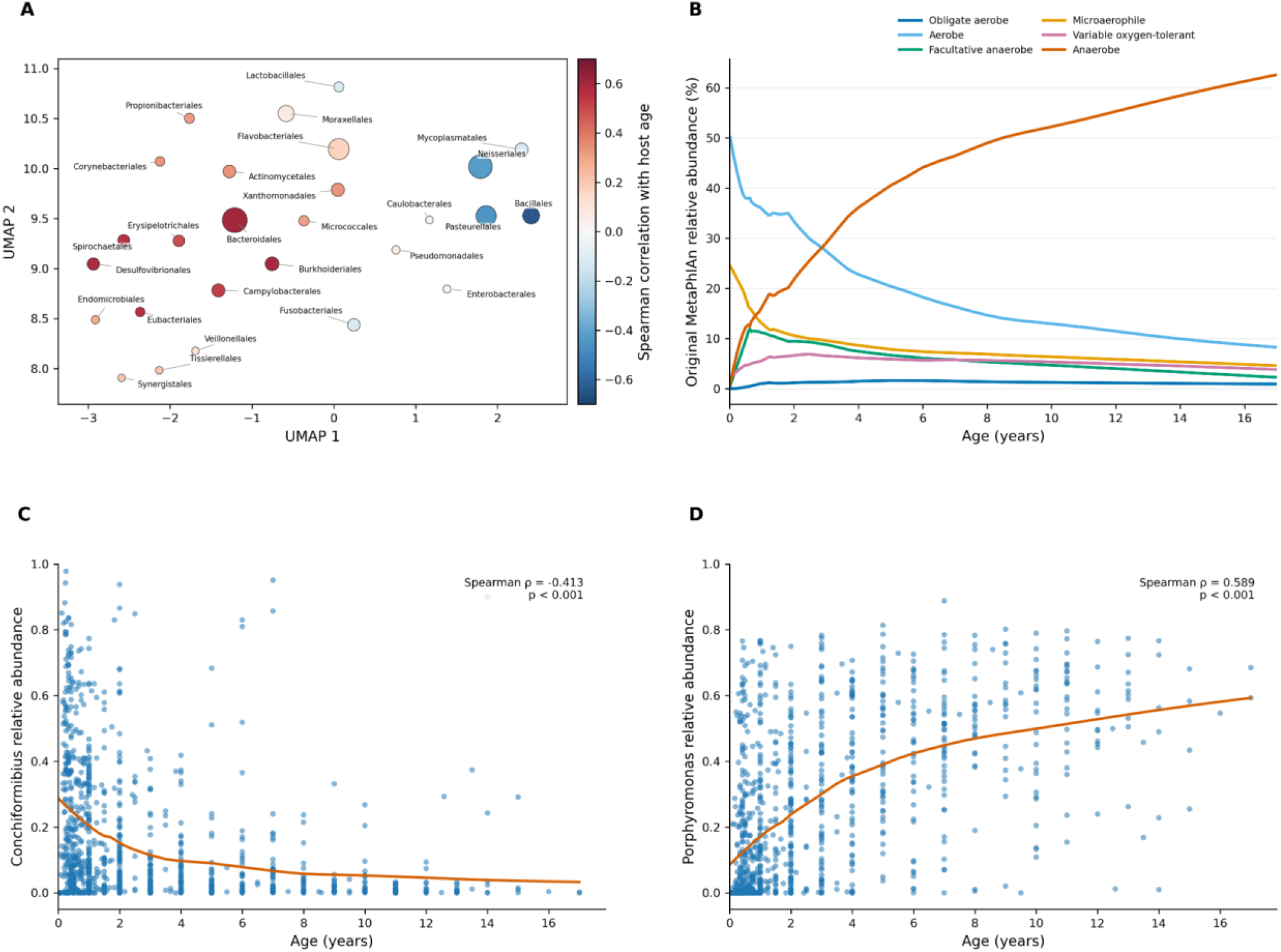
Age-associated changes in bacterial community structure, oxygen phenotype, and dominant genera. (A) UMAP projection of bacterial orders colored by Spearman correlation with host age; point size represents mean relative abundance. (B) Relative-abundance trajectories for species-level oxygen-phenotype groups across host age. (C) Relative abundance of Conchiformibius across host age. (D) Relative abundance of Porphyromonas across host age.

Species-level oxygen phenotypes were assigned as described in Methods (Figure 3B). Across individual dogs, the oxygen-tolerant fraction was highest in dogs younger than one year and declined with age (Spearman ρ = −0.532, p < 0.001), whereas anaerobic abundance increased (ρ = 0.619, p < 0.001). Across the broader 0–2-year young dog group, mean oxygen-tolerant abundance was 68.1% and mean anaerobic abundance was 14.3%. Among dogs older than 7 years, these means were 28.9% and 53.5%. The oxygen-tolerant log-ratio was also negatively associated with age (ρ = −0.615, p < 0.001). The change was already evident during the first year. The median oxygen-tolerant log-ratio decreased from 2.12 among dogs aged 0–6 months to 1.32 among dogs aged 6–12 months, corresponding to a 6.3-fold reduction (*p* < 0.001).

We focused on *Porphyromonas* and *Conchiformibius* because they were the two most abundant named genera (Supplemental Tabel S2), showed opposing age associations, and had been linked to canine age-associated oral microbiome changes in prior studies (Templeton et al., 2023; Kislik et al., 2024). Conchiformibius’s relative abundance decreased with age (ρ = −0.415, p < 0.001; Figure 3C), while Porphyromonas increased (Spearman ρ = 0.591, p < 0.001; Figure 3D), while. The full ranked genus analysis is provided in supplementary table S2.

### 4. Species-level Shannon entropy increased with age

Species-level Shannon diversity increased significantly with age (Figure 4A; Spearman ρ = 0.538, p < 0.001). Puppies (≤1 year) exhibited the lowest median diversity and greatest variability, while diversity increased in young adult dogs (>1–2 years) and remained relatively high in older age groups (>2–7 and >7 years). To examine changes during early development, we analyzed Shannon diversity across the first 12 months of life (Figure 4B). Diversity increased significantly with age during this period (Spearman ρ = 0.441, p < 0.001), with the most pronounced increase occurring during the first six months before largely plateauing. These findings suggest that much of the increase in oral microbial diversity occurs during early life, followed by more gradual changes in adulthood.

**Figure 4.**
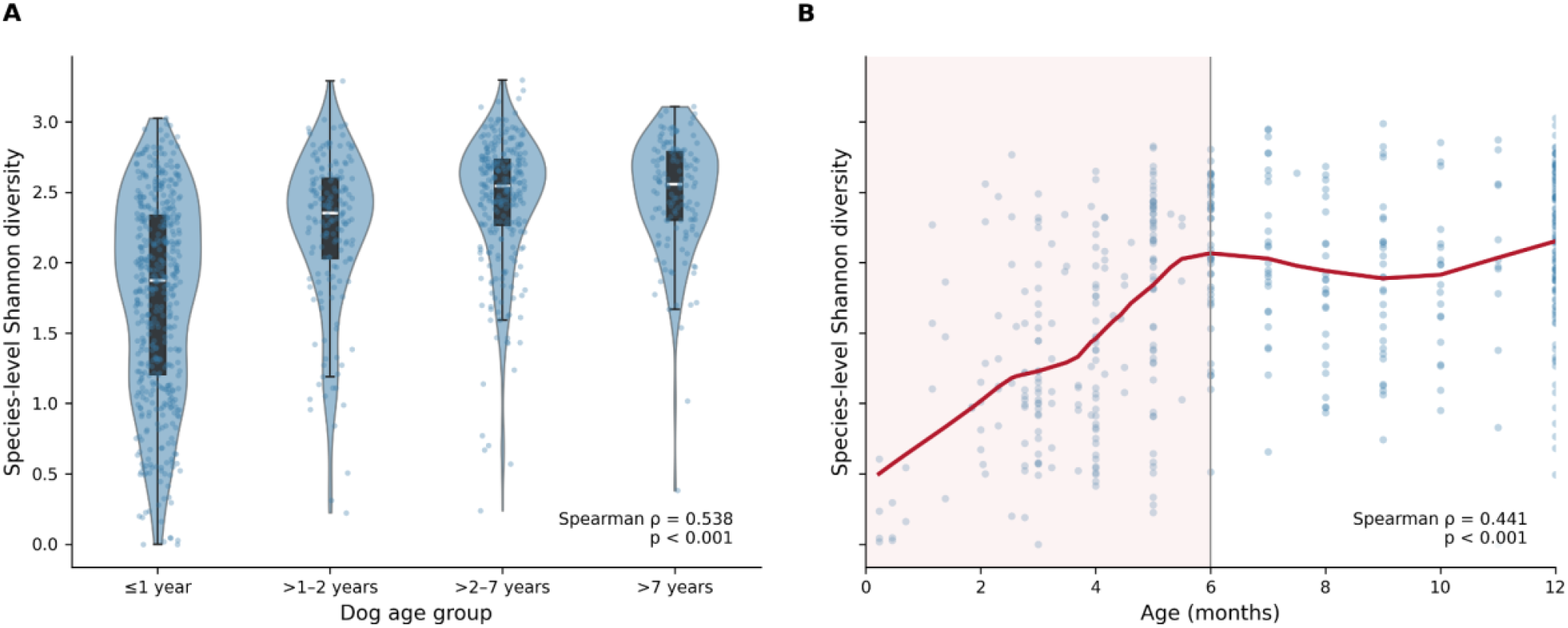
Species-level Shannon diversity across dog age groups. (A) Shannon diversity across four age groups (Spearman ρ = 0.538, p < 0.001). (B) Shannon diversity during the first 12 months of life, with a LOESS-smoothed trend (red) and the first six months shaded (Spearman ρ = 0.441, p < 0.001).

However, these diversity changes were less pronounced than the taxon-specific abundance shifts, suggesting that aging primarily alters which bacteria dominate the community rather than simply increasing overall diversity.

### 5. Age-associated restructuring of metabolic potential

To characterize the metabolic potential of the canine oral microbiome, we used HUMAnN to identify microbial gene families from shotgun metagenomic reads and map them to MetaCyc metabolic pathways. Pathway abundances reflect the representation of encoded metabolic functions within each microbial community (see Methods).

At the community level, the number of detected pathways increased with age (Spearman ρ = 0.324, p < 0.001; Figure 5A), suggesting increased functional richness in older dogs. PCA of CLR-transformed (see Methods) abundances for 301 pathways present in more than 10% of dogs identified a continuous age-associated gradient. PC1 explained 15.8% of pathway variation and correlated with age (ρ = 0.645, p < 0.001), while PC2 explained 10.1% and also correlated with age (ρ = 0.126, p = 0.002; Figure 5B). Complete PC1 and PC2 pathway loadings are provided in Supplementary Table S1.

**Figure 5.**
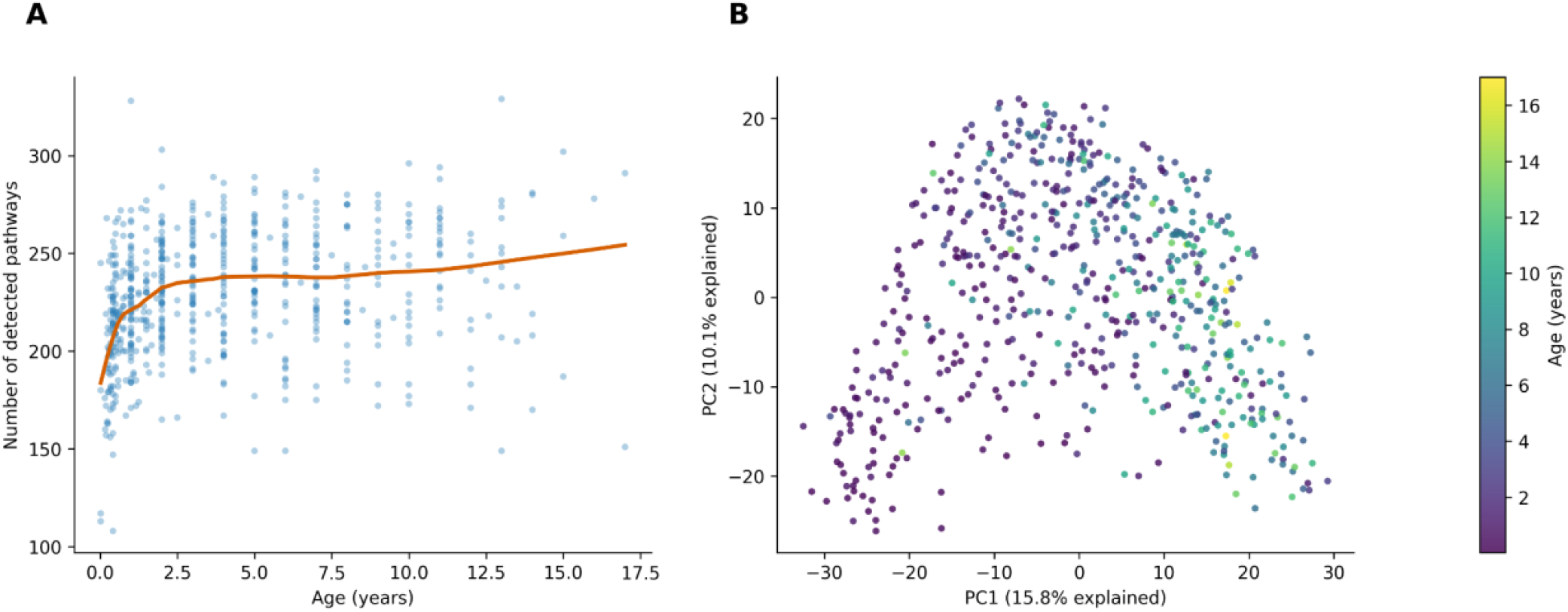
Global age associations in HUMAnN functional profiles. (A) Number of detected community-level pathways for each dog, with a LOWESS curve. (B) PCA of CLR-transformed abundances for 301 pathways detected in more than 10% of dogs; points represent individual dogs, colored by age.

Each functional-group score is the mean standardized ln(CPM + 1) abundance of pathways assigned to that group. Higher scores indicate greater relative abundance of metabolic pathways within each functional group. Amino-acid-degradation potential increased with age (Spearman ρ = 0.499, q = 2.73 × 10^−39^), as did fermentation potential (ρ = 0.250, q = 4.77 × 10^−10^). Glycolysis decreased with age (ρ = −0.381, q = 2.19 × 10^−22^), as did amino-acid-biosynthesis potential (ρ = −0.289, q = 5.34 × 10^−13^; Figure 6).

**Figure 6.**
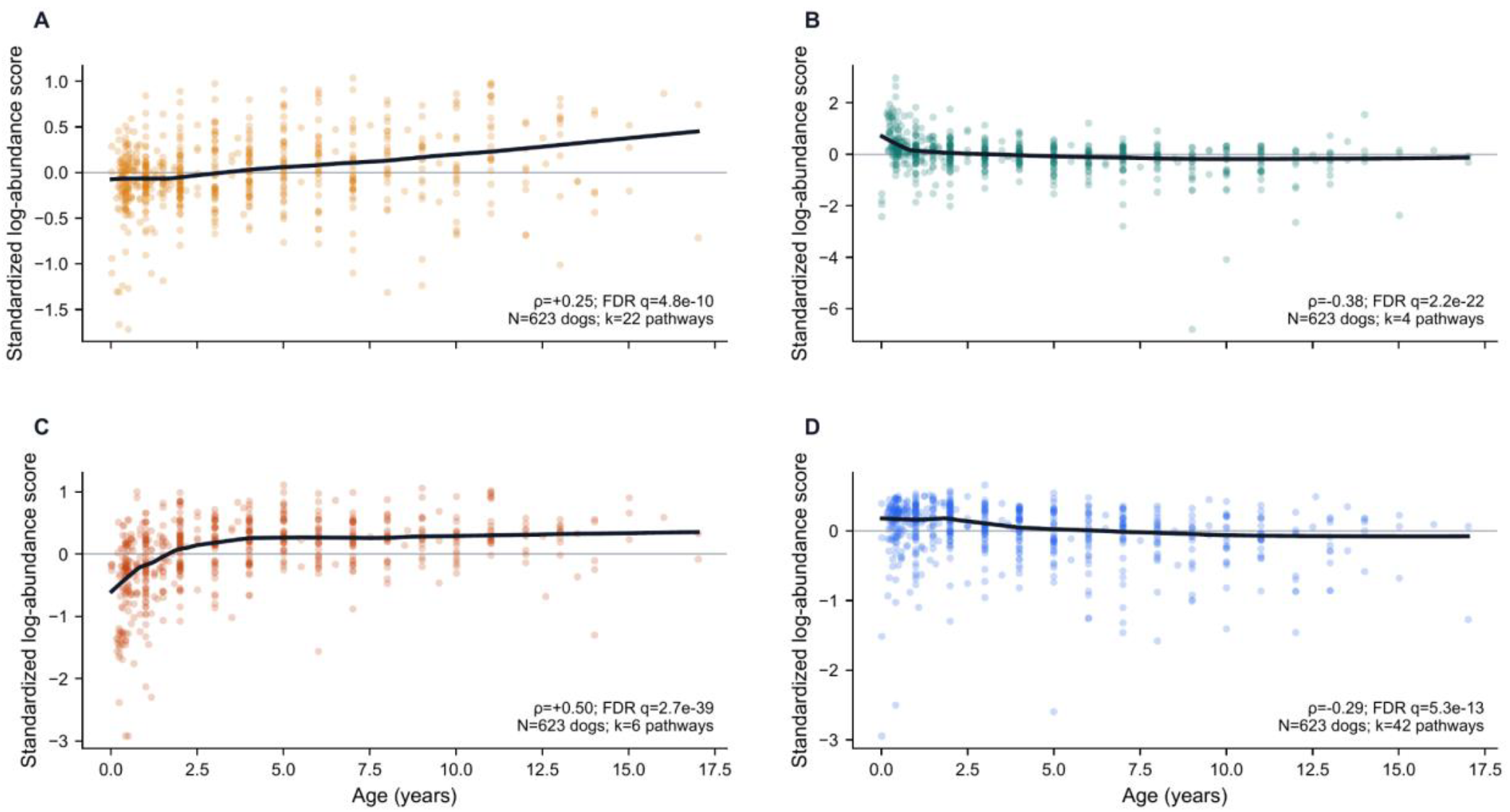
Age-associated file2meco functional-group scores. (A) Fermentation, (B) glycolysis, (C) amino-acid degradation, and (D) amino-acid biosynthesis. Points represent individual dogs (N = 695), and black lines are LOWESS fits. Panels report Spearman ρ, FDR-adjusted q, and the number of pathways in each group (k).

Respiratory pathway patterns also differed across age. Pathway detection was defined as a HUMAnN abundance greater than zero CPM. Aerobic respiration I was detected in 268 of 623 dogs, its detection not associated with age (Spearman ρ = −0.027, p = 0.499; Figure 7A). Among detected samples, abundance decreased with age (ρ = −0.332, p < 0.001; Figure 7B).

**Figure 7.**
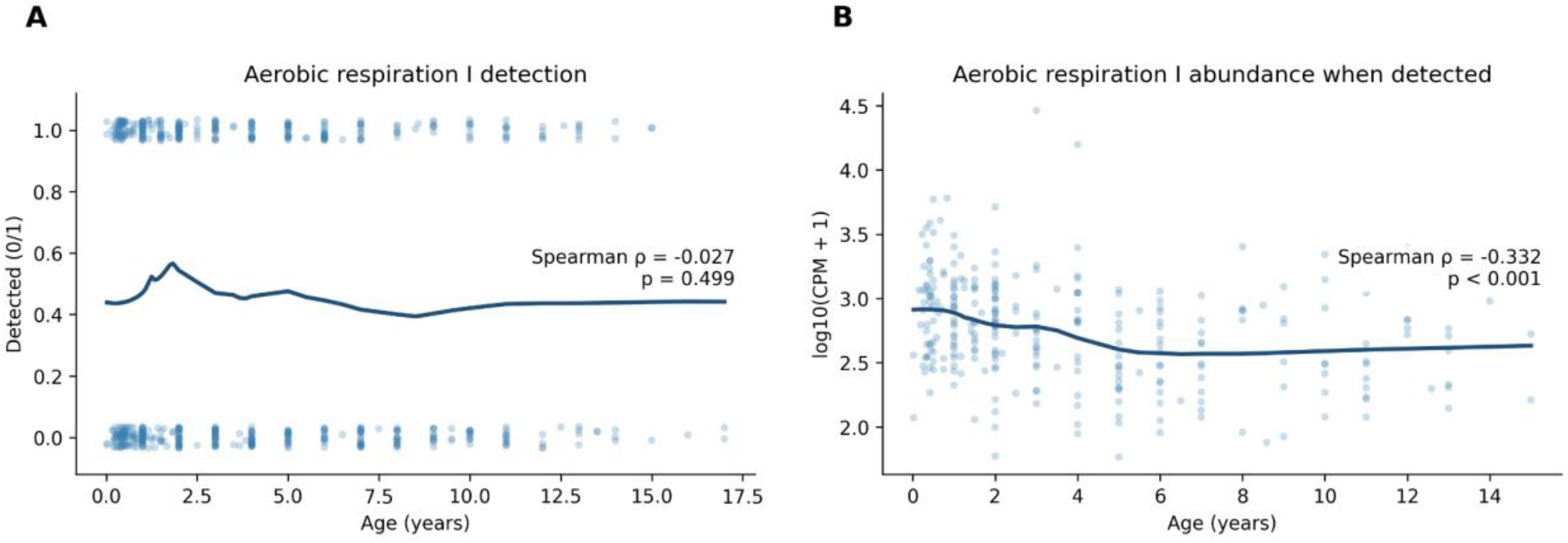
Age-associated respiratory pathway patterns. (A) Detection of aerobic respiration I across all dogs. (B) Aerobic respiration I abundance among the dogs in which it was detected.

### 6. Microbiome composition predicts host age

To assess whether oral microbiome composition could predict host age, we trained nested cross-validated Elastic Net models using 1,580 taxonomic features (see Methods). Held-out predictions were positively correlated with chronological age (Pearson r = 0.642, *p* < 0.001; R^2^ = 0.412; RMSE = 2.75 years; Figure 8). These results demonstrate that oral microbiome composition contains substantial information about host age, suggesting consistent age-associated taxonomic changes across individual dogs.

**Figure 8.**
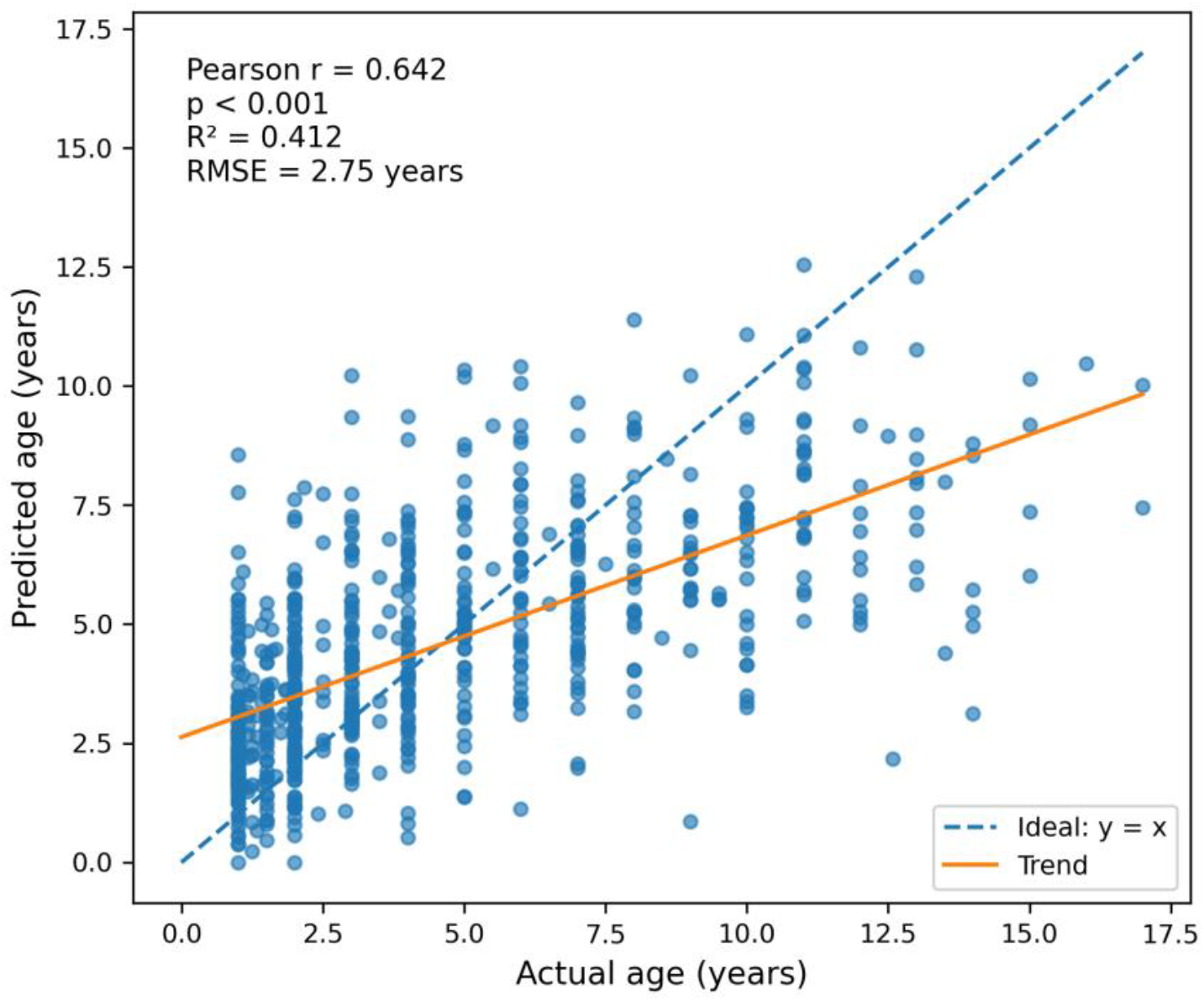
Cross-validated prediction of host age from oral microbiome composition. Points represent held-out Elastic Net predictions against chronological age. The dashed line indicates perfect agreement (y = x), and the solid line shows the fitted linear trend.

### 7. Weight-associated variation in the oral microbiome

Weight-associated changes were investigated after excluding dogs younger than one year (see Methods). In the 619-dog adult cohort with weight records, genus-level age and weight correlations were strongly inversely associated across 225 genera (Spearman ρ = −0.764, *p* < 0.001; Figure 9A). Of these, 185 taxa showed correlations of opposite sign, indicating that taxa associated with increasing age tended to show the opposite association with weight. The Elastic Net weight model showed a positive correlation between predicted and actual weight (Pearson r = 0.508, *p* < 0.001; Figure 9B). Species-level Shannon diversity also showed a weak negative association with weight group (Spearman ρ = −0.100, *p* = 0.013; Figure 9C). These results demonstrate that weight is associated with both taxonomic composition and microbial diversity.

**Figure 9.**
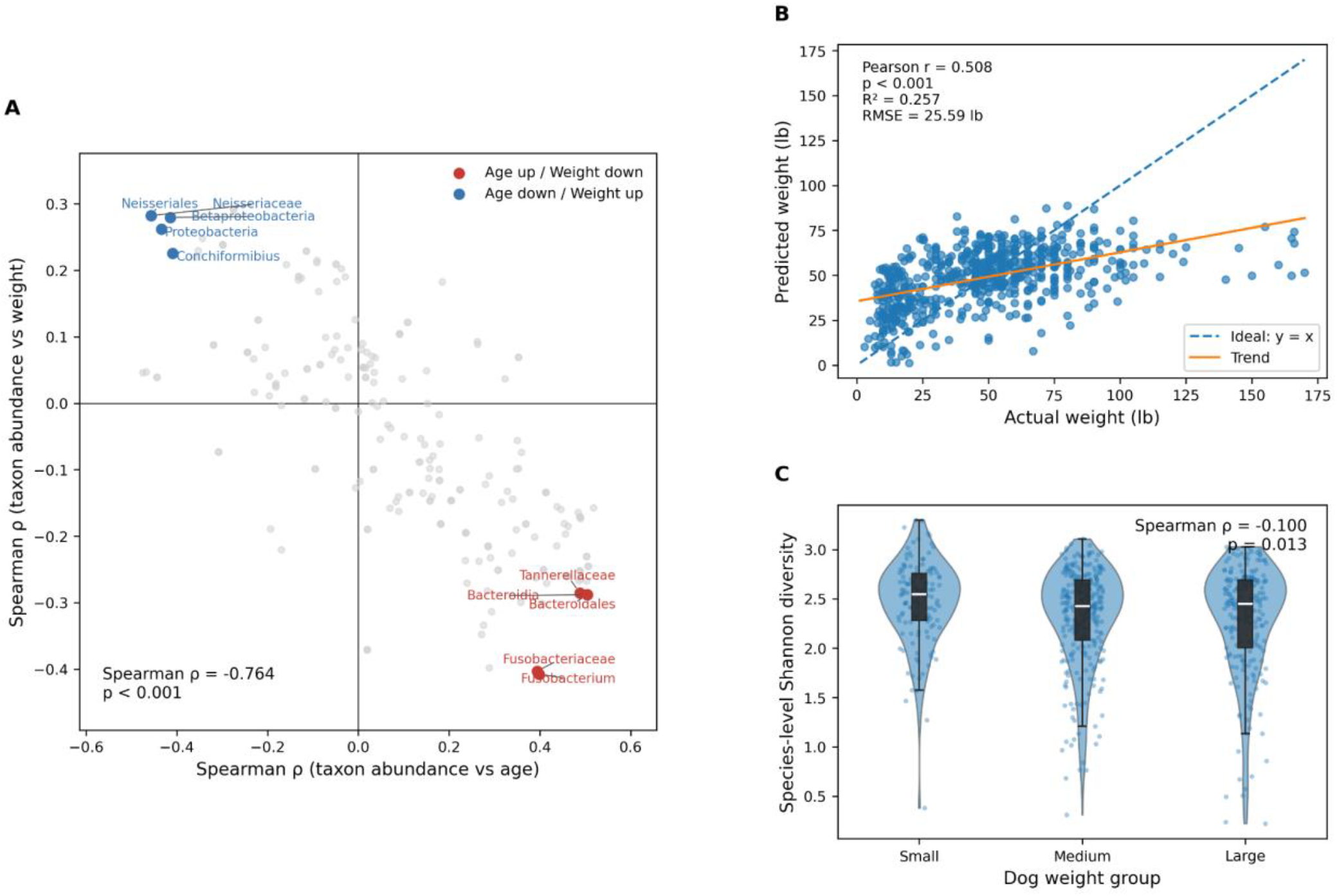
Weight-associated microbiome variation. (A) Comparison of genus-level Spearman correlations with age and weight. (B) Cross-validated Elastic Net predictions of host weight from microbiome composition. The dashed line indicates perfect agreement, and the solid line shows the fitted linear trend. (C) Species-level Shannon diversity across small, medium, and large dog weight groups.

## 4. Discussion

Age was associated with community-wide restructuring of the canine oral microbiome, characterized by a decline in oxygen-tolerant taxa, an increase in anaerobic taxa, and greater amino acid degradation potential. These changes were particularly prominent during early life (0-1 years), suggesting that the shift toward anaerobic communities begins during oral microbial community maturation. As oral biofilms develop, bacterial oxygen consumption and increasing biofilm complexity can create oxygen-limited environments that favor anaerobic growth. These conditions may also support proteolytic bacteria that utilize peptides and amino acids as nutrients. Previous work identified age-associated changes in individual canine oral taxa, particularly increased Porphyromonas and decreased Conchiformibius (Kislik et al., 2024). By jointly evaluating bacterial taxa oxygen preference with HUMAnN pathway profiles, our study extends these observations to characterize broader ecological and functional changes in the canine oral microbiome.

During early life, the oral microbiome undergoes a transition in which different bacterial groups become dominant. Tooth eruption creates new surfaces for bacterial colonization, while interactions between early colonizers promote the establishment of more complex communities (Kolenbrander et al., 2010). Morita et al. (2024) found that the oral microbiota of dogs aged 6–8 months was still developing toward adult composition. Our findings suggest that this developmental period is accompanied by a substantial change in the characteristics of the microbial community. Different sites within the canine oral cavity also support distinct microbial communities. Tooth surfaces provide stable locations for plaque formation, while the cheek and tongue promote different bacterial populations (Ruparell et al., 2020). The gingival crevice provides a more oxygen-limited environment than exposed tooth surfaces and receives protein-rich gingival crevicular fluid. These conditions can favor anaerobic bacteria capable of utilizing protein-derived nutrients (Oba et al., 2021; Marsh and Zaura, 2017). As dental biofilms mature and periodontal conditions change with age, these site-specific environments may contribute to the enrichment of anaerobic and proteolytic bacteria observed in older dogs. As biofilms become established, differences in oxygen availability and nutrient accessibility create distinct ecological niches. These conditions can support organisms with different metabolic requirements and contribute to the continued restructuring of the microbiome throughout adulthood (Marsh, 2003).

The HUMAnN analyses provide more insight about how aging may influence microbial nutrient utilization. The greater amino acid degradation potential in older dogs suggests that microbial communities become increasingly adapted to utilizing protein-derived nutrients. This greater increase may be partly explained by the increasing prevalence of periodontal inflammation with age (O’Neill et al., 2021). Inflammation increases the flow of protein-rich gingival crevicular fluid, providing additional protein substrates for bacteria in subgingival biofilms (Jakubovics, 2015; Basic and Dahlén, 2023). In dogs, Davis et al. (2016) demonstrated changes in gingival crevicular fluid proteins during progression from gingivitis to periodontitis. These conditions can favor proteolytic bacteria capable of degrading host proteins and utilizing the released peptides and amino acids. Several anaerobic oral bacteria, including *Porphyromonas*, are proteolytic and have this metabolic ability (Veillard et al., 2019). Similar metabolic patterns have been observed in human periodontal communities. Szafrański et al. (2015) reported greater carbohydrate-associated metabolism in healthy communities, while amino acid catabolism was more prominent in periodontitis. Dabdoub et al. (2016) also identified enrichment of proteolytic functions in disease-associated communities.

Together, these findings suggest that age-related changes in the periodontal environment may favor bacteria with greater amino acid degradation potential.

Respiratory pathway analyses provide another functional connection to microbial oxygen preference. The lower abundance of aerobic respiration I and greater detection of nitrate-reduction pathways in older dogs suggest a change in the microbiome’s energy-generating capabilities. Within oxygen-limited biofilm environments, bacteria capable of utilizing alternative electron acceptors can maintain energy production without relying on oxygen. Fermentation provides another means of generating energy under these conditions. These changes in respiratory potential and oxygen preference suggest that the taxonomic restructuring is accompanied by metabolic adaptation to the developing oral environment.

The increase in Shannon diversity and detected metabolic pathways in older dogs may also reflect the growing complexity of the oral microbiome. As biofilms develop, differences in local environmental conditions allow organisms with distinct physiological requirements to coexist. Interactions between these organisms can further expand the range of metabolic functions supported by the community. This provides a possible explanation for the broader functional profile and increased Shannon diversity observed in older dogs.

These findings may also help explain the relationship between age-associated microbial changes and periodontal disease. Previous canine studies have reported enrichment of anaerobic taxa in periodontal disease, particularly *Porphyromonas, Treponema*, and *Tannerella* (Davis et al., 2013; Santibáñez et al., 2021; Niemiec et al., 2022). Our findings extend these observations by identifying corresponding changes in microbial metabolic potential across age. The combination of anaerobic taxa enrichment and greater amino acid degradation potential shares the characteristics of periodontal-associated communities. This suggests that aging-related microbiome shifts may begin well before dogs reach an age at which the effects of severe periodontal disease are evident. Understanding this process could help distinguish microbial changes associated with normal development from those associated with later disease progression.

The age and weight prediction models provide further evidence that these factors are associated with microbial composition patterns. The age model’s predictive performance suggests that microbial restructuring follows a consistent pattern across dogs. Although weight was less predictive, most genus-level associations with age and weight showed opposite trends. Differences in oral anatomy between smaller and larger dogs may partially explain this pattern. Smaller dogs have proportionally larger teeth relative to their jaws, increasing the likelihood of dental crowding and plaque accumulation (Gioso et al., 2001; Wallis and Holcombe, 2020). These anatomical differences may create environments that favor anaerobic bacteria and contribute to the greater prevalence of periodontal disease in smaller breeds (O’Neill et al., 2021; Wallis et al., 2021). Previous research has also identified differences in subgingival microbial composition between small and large dogs, with smaller dogs having more gingivitis-associated bacteria, including members of Clostridiales and Erysipelotrichaceae, and lower representation of Proteobacteria associated with healthy gingiva (Wallis et al., 2021).

The Enterobacterales-enriched cluster may reflect feeding behavior. Enterobacterales are commonly found in the canine gut microbiome and their abundance can vary with diet (Tanprasertsuk et al., 2021; Bamberger et al., 2026). Coprophagy is also relatively common in dogs and may introduce fecal bacteria into the oral cavity (Hart et al., 2018).

Several limitations should be considered. This cross-sectional study cannot demonstrate longitudinal microbiome change within individual dogs. Periodontal disease status was not directly measured, limiting our ability to distinguish normal maturation from undiagnosed oral disease. Breed, diet, dental care, medication use, and household environment were not fully controlled. Differential disruption of bacterial cell walls during extraction may make some Gram-positive organisms harder to recover than Gram-negative organisms, potentially skewing final abundances. To improve the reliability of functional profiling, we restricted HUMAnN analyses to samples with at least 2 million reads. Although this reduced the influence of low sequencing depth, pathway detection may still be affected by differences in coverage, particularly for low-abundance pathways.

## Supporting information

Supplementary_Table_S1_HUMAnN_PCA_Loadings

Supplementary_Table_S2_Genus_Age_Associations

## 5. Data Availability

The entire whole-genome sequenced dataset of 1,125 dogs that was analyzed is accessible via the BioProject accession number PRJNA1525050 on the NCBI Sequence Read Archive.

